# Tuning the Mechanical Properties and Printability of Viscoelastic Skin-Derived Hydrogels for 3D Cell Culture

**DOI:** 10.1101/2025.03.12.642751

**Authors:** Estelle Palierse, Alexia Maria Mihailescu, Ira Bergquist, Cecilia Persson, Morteza Aramesh

## Abstract

In vitro investigations or tissue engineering require the creation of hierarchical and acellularized 3D structures mimicking the native environment of cells in vivo. Bioprinting provides a powerful approach to fabricating 3D architectures with precision and control. However, developing a bioink suitable for 3D cell culture remains challenging, particularly in achieving optimal rheological properties, printability and bioactivity necessary for cellular viability, functionality and growth. Here, we developed tissue-derived hydrogels with tunable gelation kinetics and rheological properties. By precisely adjusting the bioink’s physical characteristics, we optimized its printability for extrusion-based bioprinting, enabling fast fabrication of structurally stable constructs that support the formation of 3D cellular structures. A robust decellularization protocol was developed to consistently obtain porcine skin-derived dECM (decellularized extracellular matrix) hydrogels with minimal batch-to-batch variation. The influence of dECM concentration (1–5 mg/mL) on the ink’s viscoelastic properties, printability, gelation kinetics, and cellular response was investigated. Gelation kinetics varied between 7 minutes to several hours, while the storage modulus ranged between 10 to 1000 Pa. Additionally, more concentrated hydrogels led to more homogeneous prints due to their higher viscosity. Fibroblast cells infiltrated the 3D matrix of the softer hydrogels (1 and 2.5 mg/mL), forming an interconnected network. In contrast, migration was significantly restricted in the denser hydrogels (5 mg/mL). Our findings demonstrate the potential of tissue-derived hydrogels with tunable properties for 3D bioprinting applications, enabling fast and reproducible fabrication of dECM environments for cellular studies and tissue engineering, while highlighting the critical balance between mechanical and biological properties in bioink formulation.

## Introduction

Today, the demand for organ or tissue transplants is not satisfied by the number of donors, and more than 100 000 people are currently on the transplant waiting list in the US^1^. Xenotransplantation has been proposed as one of the solutions, but it is still limited by the high risk of rejection due to immunogenicity of the foreign tissue^2^. There is thus a need for a solution that enables the creation of 3D structures mimicking the 3D hierarchical native environment of human cells. Developing such constructs is also of interest for tissue modelling. Moving from 2D cultures to 3D cultures is necessary to study in vitro how cells live or react in specific environments, and crucial to move forward in the understanding of diseases or drug screening^3^. 3D cultures are also in accordance with the necessity to reduce the use of animal studies, as stated by the 3Rs policy^4^.

Bioprinting, compared to other biofabrication approaches, offers the possibility to obtain complex heterogeneous cellularized structures that can mimic the complex hierarchical 3D environment of cells in vivo.^5–11^ The more popular technique is extrusion-based bioprinting, where a bioink is loaded in a cartridge and extruded with the help of pressure, a piston or a screw.^12^ Optimal bioink’s viscoelastic properties are needed for extrudability^13^ while protecting the cells from the shear stress during printing. Additionally, sufficient stiffness and viscosity are required to avoid cell sedimentation after printing and for the shape fidelity of the printed pattern. Porosity in the gel is needed to allow for nutrient and waste transfer to and from the 3D construct after printing. Finally, bioink bioactivity is necessary for cellular viability, functionality, growth, and printability. Developing the perfect bioink is hence very challenging.

Decellularized extracellular matrix (dECM) hydrogels are of great interest for bioprinting and 3D cell culture.^11,14^ Obtained after cell removal from any tissue or organ, dECM is DNA-free, therefore little pro-inflammatory, but retains most of the biochemical components of the native ECM. The presence of growth factors and enzymes helps with cellular attachment, proliferation, and function.^15,16^ Gelation is associated to the fibrillation of the collagen present in dECM, at physiological pH and temperature, without the help of additional crosslinkers. However, despite their advantages compared to other hydrogels in terms of bioactivity, dECM hydrogels suffer from high batch-to-batch variations, slow gelation kinetics and weak mechanical properties.^14^ Their use as bioinks is therefore limited, and there is no dECM bioinks currently on the market. Despite recent progresses in the development of dECM bioinks with better printability,^11,14,17–19^ a need remains to formulate dECM hydrogels with lower batch-to-batch variation, fast gelation and the mechanical properties needed for printability.

Here we present porcine skin-derived dECM hydrogels with fast gelation and tunable viscoelastic properties. Porcine skin tissue has a composition close to the human skin and can be an accessible alternative to mimic human tissues^20^. We developed a robust protocol to obtain dECM hydrogels from porcine skin tissue, with minimal batch-to-batch variations. The influence of dECM concentration (1 to 5 mg.mL^−1^) on the hydrogel’s viscoelastic properties, printability, gelation kinetics, and cellular response was investigated. Gelation kinetics varied from 7 minutes to several hours, while storage modulus ranged from 10 Pa to 1 kPa. More concentrated hydrogels (5 mg.mL^−1^) led to more homogeneous prints due to their higher viscosity. Fibroblast cells could penetrate the 3D environment of the softer hydrogels (1 and 2.5 mg.mL^−1^), but their migration was limited in more concentrated hydrogels. Our findings demonstrate the potential of porcine-skin derived hydrogels with fast gelation and tunable viscoelastic properties for bioprinting, while capturing the critical balance between mechanical and biological properties in bioink formulation.

## Materials and methods

### Materials

Porcine skin was purchased from a local grocery store (CityGross, Uppsala, Sweden), and stored at −20°C before further use. Pepsin from porcine gastric mucosa, Papain from papaya latex, sodium dodecyl sulfate, Triton X-100, phosphate buffer saline, dimethyl methylene blue, L-cysteine, and chondroitin sulfate were purchased from Sigma-Aldrich (Solna, Sweden). Glacial acetic acid was purchased from Merck (Solna, Sweden). Mini-PROTEAN TGX gels, Laemmli sample buffer, a 10X Tris/glycine/SDS running buffer, Bio-Safe Coommassie stain and the Precision Plus Protein Standards Dual Color ladder were purchased from Bio-Rad (Solna, Sweden). Ultrapure MilliQ water (Millipore) was used for all experiments.

### Decellularization of porcine skin

Decellularization protocol was inspired and modified from published protocols.^11,21^ First, porcine skin was manually dissected to remove any visible regions of fat. The tissue was then grinded with a lab-grade grinder (Mill SM-3, Hsiang Tai) until it formed a smooth paste, and subsequently put through two 30 minute acetone washes with constant stirring to remove any trace of lipids.^22^ For all steps of the decellularization process, a ratio of approximately 10 mL of solvent per initial gram of tissue was used. The tissue was then washed with phosphate buffer saline (PBS) for 90 min, before washed with 1% Sodium Dodecyl Sulphate (SDS) in PBS for 48 h, under constant stirring. This step was followed by 45 minutes stirring in 1% Triton X-100 solution in PBS. The samples were washed for 48 h in PBS with renewal of the solution twice daily. The samples were stirred for 2 hours in a solution of 1% acetic acid and 3% hydrogen peroxide. Finally, it was washed twice with PBS for 30 min, followed by two washes with water for 15 min each. The resulting solid material was then freeze dried for 24 h. After freeze-drying, the material was grinded and washed twice with acetone for 30 min, followed by two to three washes with water to ensure removal of acetone. dECM was then freeze-dried again, grinded, and stored at −20°C before further use.

### Evaluation of decellularization

#### Biochemical characterization of native tissue and dECM

The presence of the major skin components in dECM, i.e., collagen, elastin and sulfated glycosaminoglycans (GAGs), as well as the content of DNA, were measured by biochemical assays, and compared with native tissue. Collagen contents were measured using the Sircol Soluble Collagen Assay (Biocolor). Samples were pre-treated by 16 h of pepsin digestion in 0.5 M acetic acid at 4 °C at a concentration of 5 mg.mL^−1^ for dECM and 20 mg.mL^−1^ for tissue, and then assessed according to the kit instructions. Elastin contents were measured using the Fastin Elastin Assay (Biocolor), according to the manufacturer instructions. For GAGs and DNA quantification, dECM and tissues were first digested in a papain solution (125 mg.ml^−1^ papain in 0.1 M sodium phosphate with 5 mM Na_2-_EDTA and 5 mM cysteine-HCl at pH 6.5), at a concentration of 50 mg.mL^−1^ for 16 h at 60 °C. The solution was then centrifuged 10 min at 13 000g, and the supernatant was filtered with a 0.22 µm filter. Sulfated GAG content was assessed using the dimethyl methylene blue (DMMB) method.^23^ The DMMB reagent solution consisted of 38 µM DMMB, 40 mM glycine and 27 mM NaCl in 10 mM acetic acid. A standard curve consisting of chondroitin sulphate (0-250 µg/mL) in MilliQ water was used. 20 µL of the standards and samples were loaded in triplicates onto a 96-well plate followed by 200 µL of the DMMB solution. The plate was shaken for 1 min in the dark and then immediately measured for absorbance at 525 nm. The DNA content was measured using Invitrogen’s Quant-iT PicoGreen dsDNA assay kit according to the manufacturer’s instructions. For all the tests, the absorbance or fluorescence of the solutions were measured using a Tecan Infinite M200 plate reader. All tests were performed in triplicates.

#### Visualization of native tissue and dECM

Native tissue and dECM gels were observed with confocal imaging. Porcine skin specimens for imaging were prepared from the frozen tissue by slicing 10 μm sections in a cryostat microtome (Micron HM560, Germany). Before staining, sections were thawed at room temperature for 1 h and fixed with 4% PFA for 15 min. dECM gels at a concentration of 5 mg.mL^−1^ were obtained after 24 h incubation of pre-gels at 37°C, as described in the next section, and were fixed with 4% PFA for 15 min. Before staining, both native tissue sections and dECM gels were permeabilized with 0.1% Triton X-100 in PBS for 10 min, before being washed three times with PBS. They were incubated with 2% BSA in PBS for at least 1 h at room temperature. For staining, they were incubated with Actin staining 488 (2 drops/mL) and DAPI (5 µg/mL) for 1 h at room temperature in the dark.

Confocal imaging was performed with a SP8 confocal laser scanning microscope (Leica), using a 10 × air objective. Images were acquired with a 1024 × 1024 resolution with 400 speed.

### Preparation of dECM pre-gels and gels

dECM pre-gels were obtained after the enzymatic digestion of freeze-dried dECM at a concentration of 5 mg.mL^−1^, with pepsin (2 mg of pepsin per 100 mg dECM) at 4 °C in a 0.5 M solution of acetic acid (protocol adapted from Voytik-Harbin et al^24^). After solubilization, the pH was adjusted to 7.2−7.4 using sodium hydroxide at 10 and 1 mol.L^−1^ and acetic acid at 1 and 0.5 mol.L^−1^, while keeping the temperature below 10 °C. The solution was stirred for 1 h at 4 °C to let the pH stabilize. The pH was readjusted if necessary. The solution was then centrifuged at 1000 ×g for 5 min to remove any non-dissolved lipids bits. To prepare pre-gels at lower concentrations of 2.5 and 1 mg.mL^−1^, pre-gels were diluted with PBS at pH 7.4. Pre-gels were stored at 4 °C and used or studied within a week.

The dECM gels at 5 mg.mL^−1^ and 2.5 mg.mL^−1^ concentrations were obtained after 24 h incubation of pre-gels at 37 °C. 1 mL of pre-gel was cast in a 20 mm-diameter Teflon mold and kept under wet conditions during incubation.

### Rheological characterization

Rheological investigations were conducted with a Rheometer HR10 (Discovery Series, TA Instruments – Waters AB, Solna, Sweden) in parallel plate mode using a 20 mm diameter geometry and Peltier plate. Before the start of each test, the sample was agitated with a vortex for 30 s to homogenize the solution, and 350 µL of pre-gel was loaded onto the plate. Silicone oil AR 20 (SigmaAldrich) was applied around the sample, to avoid drying during experiments.

The viscosity of the pre-gels was measured with flow sweep measurements at 4 °C, with a shear rate from 0.1 to 1000 s^−1^. Gelation kinetics was studied with time sweep measurements, performed at 1 Hz and 1% shear, at 4 °C for 300 s and 1 h at 37 °C. Gelation time was extracted for when the phase angle decreased and reached a value below 10°. Strain sweep measurements 37 °C were performed on dECM gels with 20 mm diameter at 5 mg.mL^−1^ and 2.5 mg.mL^−1^ concentration, at 1 Hz frequency, with a stress ranging from 0.1 to 100%.

### Turbidimetric gelation kinetics

The absorption of 200 µL of pre-gel solutions was recorded over 4 h at 405 nm, at 37 °C, with a Tecan Infinite M200 plate reader. Absorption was recorded every 5 min and the plate was agitated before each measurement for 1 min. Absorbance values were normalized by the absorbance of a PBS solution.

### Observation of microstructure

The microstructure of the gels at 1, 2.5 and 5 mg.mL^−1^ dECM concentration was observed by scanning electron microscopy (SEM). For that, pre-gels were casted and let to gel overnight at 37 °C, to obtain gels with a thickness of roughly 5 mm. These gels were frozen with liquid nitrogen and directly freeze-dried for 30 h. Dried gels were then carefully broken to expose the inner part of the gel. Samples were carefully mounted on a carbon tape, with the inner part facing upwards. The samples were imaged with a ZEISS Crossbeam 550 FIB/SEM, with an accelerating voltage of 0.5 or 1 kV, and a current of 100 or 200 pA, at a working distance of 2 mm. No metallic coating was necessary for these observations.

### Printability tests

An open source bioprinter based on the E3D motion system was used for extrusion-based printing.^25^ Pre-gels at 1, 2.5 and 5 mg.mL^−1^ were extruded through a 30G syringe with an inner diameter of 150 µm. Printing occurred in air at 20 °C, at 300, 600 and 1000 mm.min^−1^. The temperature of the pre-gels was kept at below 15 °C, to avoid gelation in the cartridge, and the plate was at room temperature. A one-layer serpentine was printed on a glass slide to determine resolution and spreading of the printed construct by measuring the width of the extruded filament.

In a second set of experiments, the same one-layer serpentine was printed on a polystyrene petri dish (Sigma Aldrich), at 600 mm.min^−1^, with pre-gels at 2.5 and 5 mg.mL^−1^. Finally, to test for the layer stacking, a square plate with the dimension 10×10×0.15 mm^3^ was printed with the following parameters: 2 or 4 layers (height = 0.3 mm or 0.6 mm), infill grid 20% with 1 wall, 0 top, 0 bottom.

Pictures were taken directly after printing with a digital microscope (Dino-Lite Europe, Almere, The Netherlands), using the Dinocapture 3.0 software. The images were analyzed with ImageJ software afterwards. The width of the printed material for the different concentrations and printing speeds were measured at eight locations for each print.

### In vitro cell culture

L929 murine fibroblasts (ECACC 85011425) were grown in Dulbecco modified eagle medium (DMEM) with high glucose and L-glutamine (Gibco 11965092) and supplemented with 10% Fetal Bovine Serum (FBS) (Gibco 11560636) and 1% penicillin/streptomycin (Sigma-Aldrich, P4333), at 37 °C with 5% CO_2._ The cells were detached from culture flasks, using TrypLE (Gibco, 11528856) for 5 minutes at 37 °C, and were resuspended to a concentration of 34 000 cells.mL^−1^, prior to culturing on top of dECM gels.

For cell culture experiments, dECM pre-gels at 5 mg.mL^−1^ were diluted with 10X PBS to adjust for the osmotic pressure in the pre-gels. To prepare 5 mL of adjusted pre-gel, 500 µL of 10X PBS was added to 4.5 mL of pre-gel. This adjusted pre-gel was then diluted with 1X PBS to obtain pre-gels at 2.5 and 1 mg.mL^−1^.

#### Study of cell proliferation on top of dECM gels

To study the proliferation of fibroblasts on top of dECM gels, 40 µL of pre-gel was cast in the wells of 96 well plates, and incubated overnight for gelation, at 37 °C, 90 % humidity and 5 % CO_2._ After incubation, 100 µL of fibroblasts cell suspension were seeded on top of the gel to reach a cell density of 10^4^ cells.cm^−2^. Cells were cultured up to 7 days and fixed after 24 h, 72 h and 7 days. Cell medium was changed after 72 h.

### Immunostaining

Samples were fixed with 4% paraformaldehyde in PBS (PFA, Sigma Aldrich) for 10 min at room temperature, and washed with PBS three times. They were stored in PBS at 4 °C for up to 2 weeks before immunostaining. First, they were permeabilized with 0.1% Triton X-100 in PBS for 10 min, before being washed three times with PBS. Then, they were incubated with 2% BSA in PBS for at least 1 h at room temperature. For staining, they were incubated with Actin staining 488 (2 drops/mL), a-Tubulin AF-647 (5 µg/mL) and DAPI (5 µg/mL) for 1 h at room temperature in the dark. After three washes in PBS, they were stored at 4°C covered in PBS before imaging.

### Confocal fluorescence microscopy imaging

Confocal fluorescence microscope imaging was performed with a SP8 confocal laser scanning microscope (Leica), using a 10 × air objective. Images were acquired with a 1024 × 1024 pixels resolution with 400 lines/sec speed.

### Statistical analysis

OriginLab 2019 software was used to conduct statistical analysis. All results except the decellularization results were analysed using a one-way analysis of variance (ANOVA) test and differences between groups were determined with Tukeýs post hoc analysis. Decellularization results were analysed using unpaired student t-test. A p-value < 0.05 was considered statistically significant.

## Results and discussion

### Decellularization

Unlike other studies focused on the dermal layer,^26,27^ here we aimed to use the whole skin tissue without isolating epidermis, dermis and hypodermis. However, as the skin source was derived from the food industry, the initial tissue contained a significant amount of fat. The first step of decellularization therefore involved removal of the fat lipids. The visible parts were cut and removed manually before washing the tissue several times with acetone to remove the remaining lipids. Then, the decellularization process was conducted by successive washing steps with surfactants or oxidant solution, with extended washing steps in PBS in between the steps. An additional washing step with acetone was conducted if lipids were still present within the decellularized solution. The efficiency of lipid removal can be seen by measuring the concentration of major ECM components, *i.e.,* collagen, sulfated GAGs and elastin, in µg per mg of dry tissue or ECM, presented in Fig. 1a. Protein and glycan concentrations measured in the native tissue were largely variable, due to inherent inhomogeneity within the tissue. Protein and glycan contents were very low in the native tissue, with only 63 µg.mg^−1^ of collagen, 1.2 µg.mg^−1^ of sulfated GAGs and 5.8 µg.mg^−1^ of elastin. On the contrary, the results were very homogeneous for dECM. Depending on the component, the concentration in dECM was more than 10 times higher than in native tissue, which confirms that dECM is composed mainly by its major structural proteins and glycans, but not lipids. The preservation of the ECM components after decellularization can be assessed by presenting the latter results in percentage of the initial quantities in the native tissue used for decellularization (Fig. 1b). The quantity of collagen and elastin are preserved in dECM, but sulfated GAGs are lost significantly after decellularization. However, sulfated GAGs are usually found difficult to preserve during decellularization.^26,28^

**Fig. 1.**
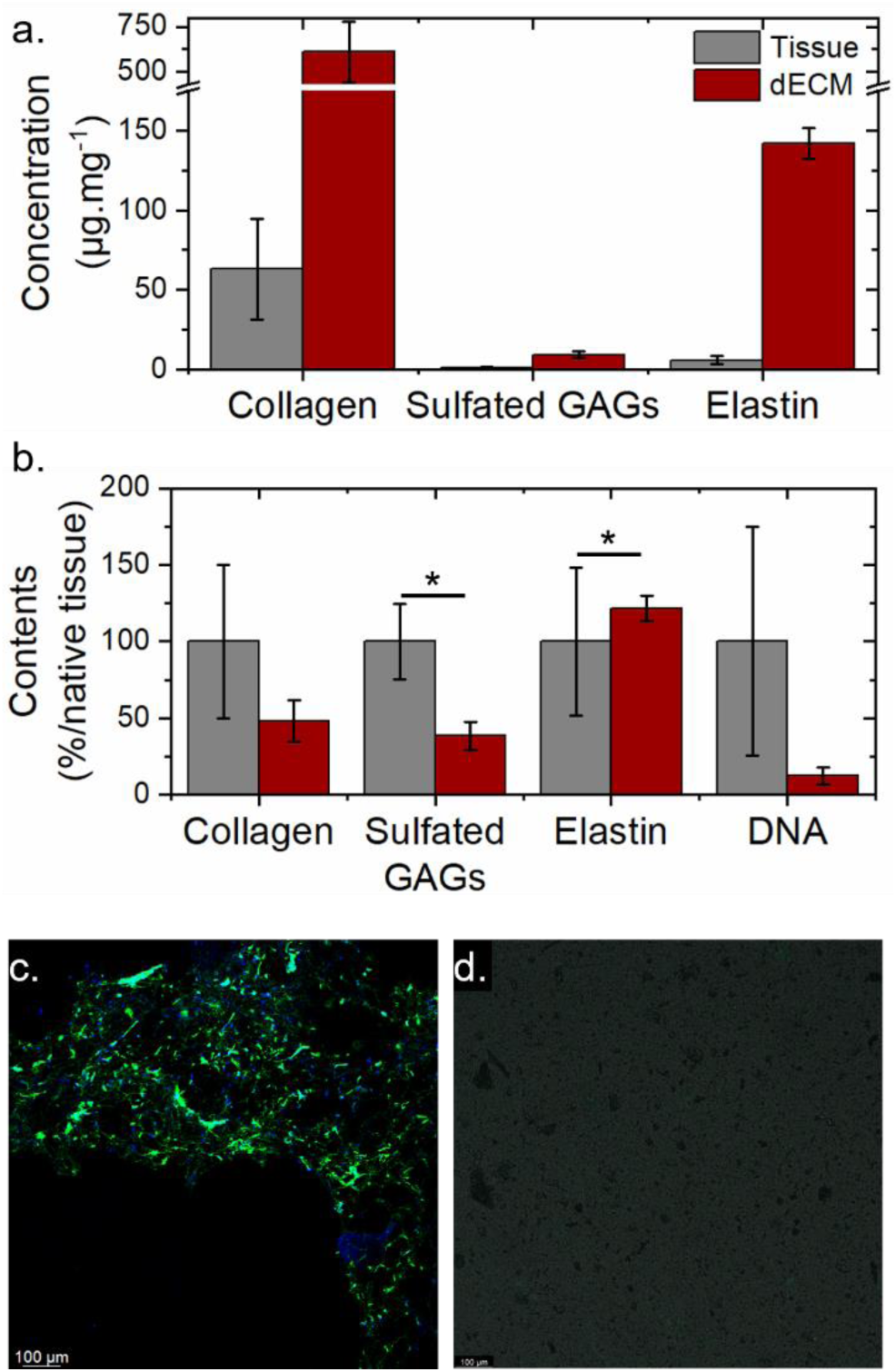
Biochemical and microscopical analysis of decellularized tissues. a. ECM component concentration presented in µg per mg of dry tissue or dry dECM. b. Maintenance of ECM components (collagen, sulfated GAGs, elastin) and DNA after decellularization. c,d, Confocal microscopy imaging, comparing cellular content before (native tissue) and after decellularization (dECM hydrogel), respectively. DAPI (blue) and F-actin (green) staining indicate the presence of DNA and actin (representing intracellular proteins) All experiments were performed in triplicates. Error bars represent s.d.

The removal of DNA is crucial for efficient decellularization. We observed a reduction of DNA in dECM compared to native tissue, with ∼12% of DNA present after decellularization, without using any DNase in the process. To confirm the efficiency of decellularization, sections of native tissue and dECM gels were observed with confocal imaging after DAPI and F-actin staining (Fig. 1c-d). Native tissue sections (Fig. 1c) showed the presence of cells labelled by DAPI, whereas the fluorescence signal is absent in dECM gels (Fig. 1d). Similarly, F-actin fluorescence was present in the native tissue sections but absent in dECM gels.

### dECM pre-gel and hydrogel

dECM was solubilized by an enzymatic digestion with pepsin for 24 h at acidic pH. dECM pre-gels were obtained after adjusting the pH of the dECM solution to 7.2 – 7.4. Here, dECM was solubilized at a concentration of 5 mg.mL^−1^. After dilutions with PBS, pre-gels at 2.5 and 1 mg.mL^−1^ were obtained. The viscosity of the pre-gels was assessed at 4 °C, as a function of pre-gel concentration, through flow sweep measurements. Fig. 2a shows the low viscosity of the pre-gels, which decreases with decreasing concentration. At 10 s^−1^ shear rate, dECM pre-gels have a viscosity of 0.16 Pa.s at 5 mg.mL^−1^, 0.02 Pa.s at 2.5 mg.mL^−1^ and 0.006 at Pa.s at 1 mg.mL^−1^. All pre-gels possess shear-thinning properties with lower viscosity at higher shear rate. The range of viscosity of dECM pre-gels, combined with their shear thinning properties, make them suitable for injectability or extrudability of pre-gels.^29^

**Fig. 2.**
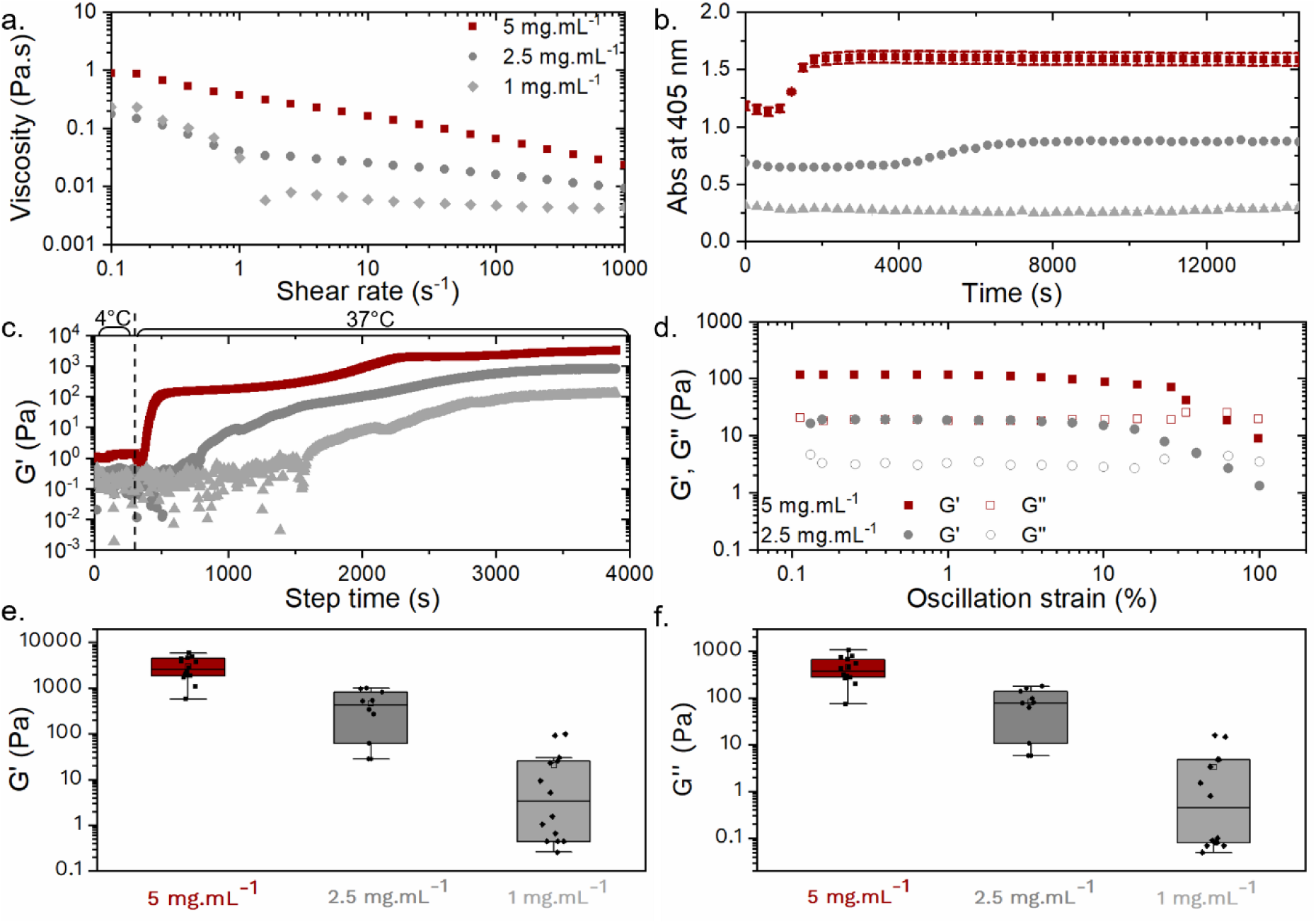
Rheological characterization of dECM pre-gels and hydrogels at different concentrations (5, 2.5 and 1 mg.mL^−1^). a. Viscosity of the three pre-gels measured at 4°C. b. Gelation kinetics assessed by turbidimetry, measuring absorbance at 405 nm over time at 37 °C. c. Gelation kinetics assessed by a rheometer, measuring storage modulus over time at 37°C. d. Shear thinning properties of dECM gels precast at 37°C. e.,f Storage (G’) and loss (G**’’**) moduli of the corresponding dECM hydrogels after gelation, measured by a rheometer. Each dot corresponds to one replicate.

The gelation kinetics of dECM hydrogels were assessed by two complementary techniques: turbidimetry and rheology, as seen in Fig. 2b and Fig. 2c, respectively. With turbidimetry measurements, the absorbance of pre-gel solutions was monitored over time at 37 °C. The increase in absorbance is associated to the gelation of dECM.^27^ At 5 mg.mL^−1^, the absorbance started to increase after 15 min and the plateau was achieved after 1 h. At 2.5 mg.mL^−1^, the absorbance increased after 1 h and the plateau was achieved after 2.5 h. Nevertheless, the gelation was not detectable with this technique at 1 mg.mL^−1^. With rheology measurements, the storage and loss moduli of dECM pre-gels were monitored over time. For the first 5 min, temperature was kept at 4 °C, before being raised to 37 °C. The temperature reached 37 °C within 1 min. By increasing the temperature, storage and loss moduli increased and the storage modulus curve crossed the loss modulus, indicating the gelation (Fig. S2). The crossing occurred within 1 min at 5 mg.mL^−1^. The gelation time was defined as the time when the phase angle reached its minimum. This time was 6.8 min, 22.5 and 36 min for 5, 2.5 and 1 mg.mL^−1^ pre-gels, respectively. The gelation was found to be faster when studied by rheology compared to turbidimetry. This is due to a better conduction of temperature on the rheometer plate compared to inside the plate-reader. The heating is very local on the Peltier plate, which is not the case inside the plate-reader. Overall, our results indicate that gelation kinetics varies with pre-gel concentration, as expected. The gelation time with dECM pre-gels at 5 mg.mL^−1^ is remarkably fast, compared to what has already been reported in the literature for porcine-skin derived hydrogels.^26,27^

Interestingly, for each concentration, time sweep curves are characterized by two phases. At 5 mg.mL^−1^, a first plateau reaches at 100 Pa, followed by a second at 3 kPa. At 2.5 mg.mL^−1^, plateau are at 50 Pa and 500 Pa, respectively, and at 10 and 100 Pa for 1 mg.mL^−1^ gels. Two plateau curves are not common for collagen or dECM gels, but have been observed in the case of low concentrated gels.^30,31^ The value of the storage modulus of the first plateau corresponds to the value of the storage modulus of gels when molded and prepared overnight at 37 °C (Fig. 2d). When prepared under these conditions, the hydrogels are completely saturated with water. The first plateau may then correspond to swollen hydrogels and the second plateau to the dry form of the hydrogel. This is further confirmed by observing correlated patterns between the axial force and the phase angle during the gelation kinetics (Fig. S2d,e). Indeed, a large negative peak in axial force occurred just before the second plateau, while the phase angle remained constant. A negative peak in axial force during rheology studies is associated with drying of the material, the force going down due to the contraction of the material.

The final storage and loss moduli after gelation kinetics are presented in Fig. 2e, for the three concentrations. It is obvious that the dispersity of the values at 5 mg.mL^−1^ is very small, measured for different batches. This is an indication of high consistency across the batches for the obtained pre-gels. However, dilutions of pre-gels with PBS leads to more variability in G’ values, especially at 1 mg.mL^−1^, where G’ varies from 0.1 to 100 Pa. This could be associated with a higher sensitivity of the gelation process at low concentrations to the environmental conditions during pre-gel preparation (temperature, humidity). In this study, dECM hydrogels at 5 mg.mL^−1^ reach the same storage modulus values as gels extracted from harder tissues like tendons,^32^ which are generally stiffer. Another advantage of the hydrogels presented here is their relatively low concentration. Hydrogels are developed here between 1 and 5 mg.mL^−1^, corresponding to 0.1 to 0.5% w/v. These concentrations are in the lower range of what it is commonly used for dECM hydrogels, where it is not rare to formulate gels with concentrations of up to 10%.^11,27,32,33^ The high storage moduli achieved in this work can be attributed to a lower degree of dECM digestion, minimizing fragmentation of the native proteins.^21,34^ Indeed, in this work, we decreased the pepsin concentration to 2 mg per 100 mg of dECM, and digestion occurred at 4 °C for only 24 h. In the literature, digestion time is usually between 24 and 72 h, and room temperature is commonly used, as has been previously reviewed by Saldin et al.^35^ The molecular weight of the collagen macromolecules present in dECM pre-gel was measured by SDS page assay and compared with standard collagen from rat tail tendon (Fig. S1). Macromolecules in dECM are larger than 150 kDa, which confirms that they were not extensively fragmentized during digestion.

The microstructure of the gels was observed with scanning electron microscopy (Fig, 3). For that, dECM gels were freeze-dried before imaging. At small magnification, the freeze-dried gels at 5 and 2.5 mg.mL^−1^ present a very organized structure, with two dimensional sheets linked by single fibers (Fig. 3a,b for 5 mg.mL^−1^ and Fig. 3d,e for 2.5 mg.mL^−1^). At higher magnification, we can observe that the sheets are composed by fibers that are randomly oriented (Fig. 3c for 5 mg.mL^−1^ and Fig. 3f for 2.5 mg.mL^−1^). The structure is more dense at 5 mg.mL^−1^ compared to the gel at 2.5 mg.mL^−1^. The gel at 1 mg.mL^−1^ present the same organized structure (Fig. 3g-i). The number of fibers is much lower compared to the more concentrated gels. In addition to the fibers, the images of the gels at 2.5 mg.mL^−1^ and 1 mg.mL^−1^ present some cubical structures that become the main component at 1 mg.mL^−1^. These are associated to the NaCl and phosphate salts present in the PBS, used to dilute the pre-gels at the right concentration. With freeze-drying, the salts are still presents in the gels. It is worth to note that these images represent the microstructure of the freeze-dried gels, which can be different to the gel in their wet state.

**Fig. 3.**
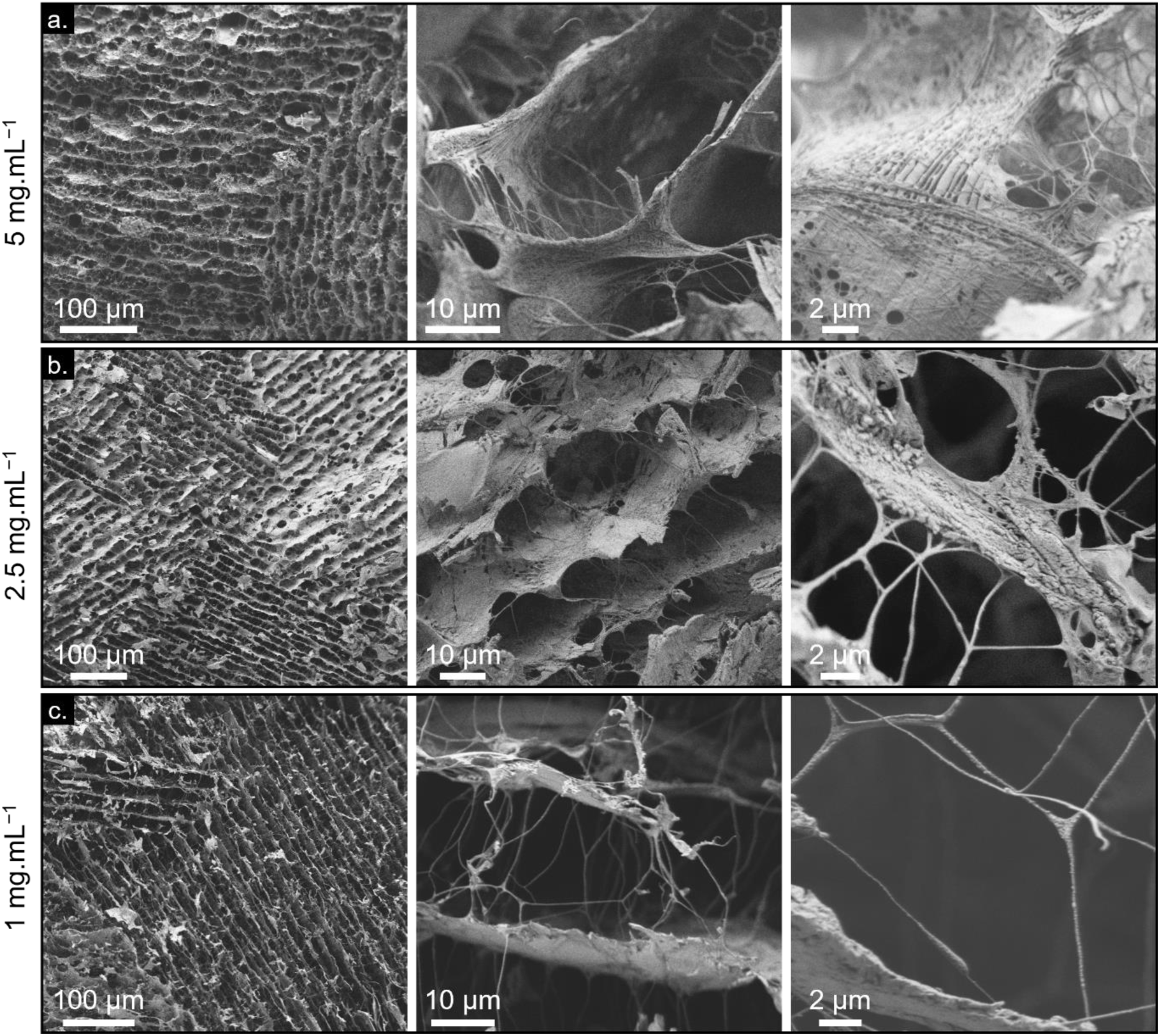
Scanning electron microscopy images of freeze dried dECM gels. a. 5 mg.mL^−1^, b. 2.5 mg.mL^−1^, c. 1 mg.mL^−1^

### Printability

The printability of the dECM pre-gels at the three concentrations was studied with extrusion-based 3D printing. In a first study, a serpentine pattern consisting of one layer was printed at three different speeds: 300, 600 and 1000 mm.min^−1^ on a glass slide to determine the best parameters for printing. A needle of 30G, corresponding to an inner diameter of 150 µm was used for all the studies. Shape fidelity of the printed constructs was assessed by comparing the diameter of the needle and the width of the filament directly after printing. After-print pictures were taken with a digital microscope and the thickness of the filament was measured at 8 different points. Representative after-print pictures at 600 mm.min^−1^ are presented in Fig. 4a-c, and in Supplementary Information for the other printing speeds (Fig. S3). For all dECM concentrations and printing speeds the serpentine shape is respected, except for the smallest gap of 1 mm. Here, filaments fused because of high spreading on the glass slide. Spreading of the filament is quantified by measuring its width directly after printing and presented in Fig. 4d. For all concentrations and speeds, the width was between 400 and 800 µm, which is ∼3 times larger than needle’s inner diameter. The shape fidelity is presented as the ratio DS/DF, where DS is the inner diameter of the syringe and DF the diameter (width) of the filament after printing measured on the glass slide. DS/DN are presented in Fig. 4e. For all concentrations and speeds, DS/DF was between 0.2 and 0.25. There is no statistical difference in the mean of the filament diameter between the concentrations for each speed and between the speeds for each concentration. This absence of statistical difference can be attributed to the high spreading of the inks on the glass slides, due to the hydrophilicity of glass, and in general low viscosity of the ink at each concentration. However, not all the prints have the same homogeneity. The standard deviation to the mean of the width of the printed filament for all the speeds and the concentrations is plotted in Fig. 4d. We observed that there is less variability for the prints at 5 mg.mL^−1^, and more variability for the prints at 1 mg.mL^−1^, with an intermediate variability at 2.5 mg.mL^−1^. In between the speeds, 600 mm.min^−1^ is the speed that leads to higher shape fidelity.

**Fig. 4.**
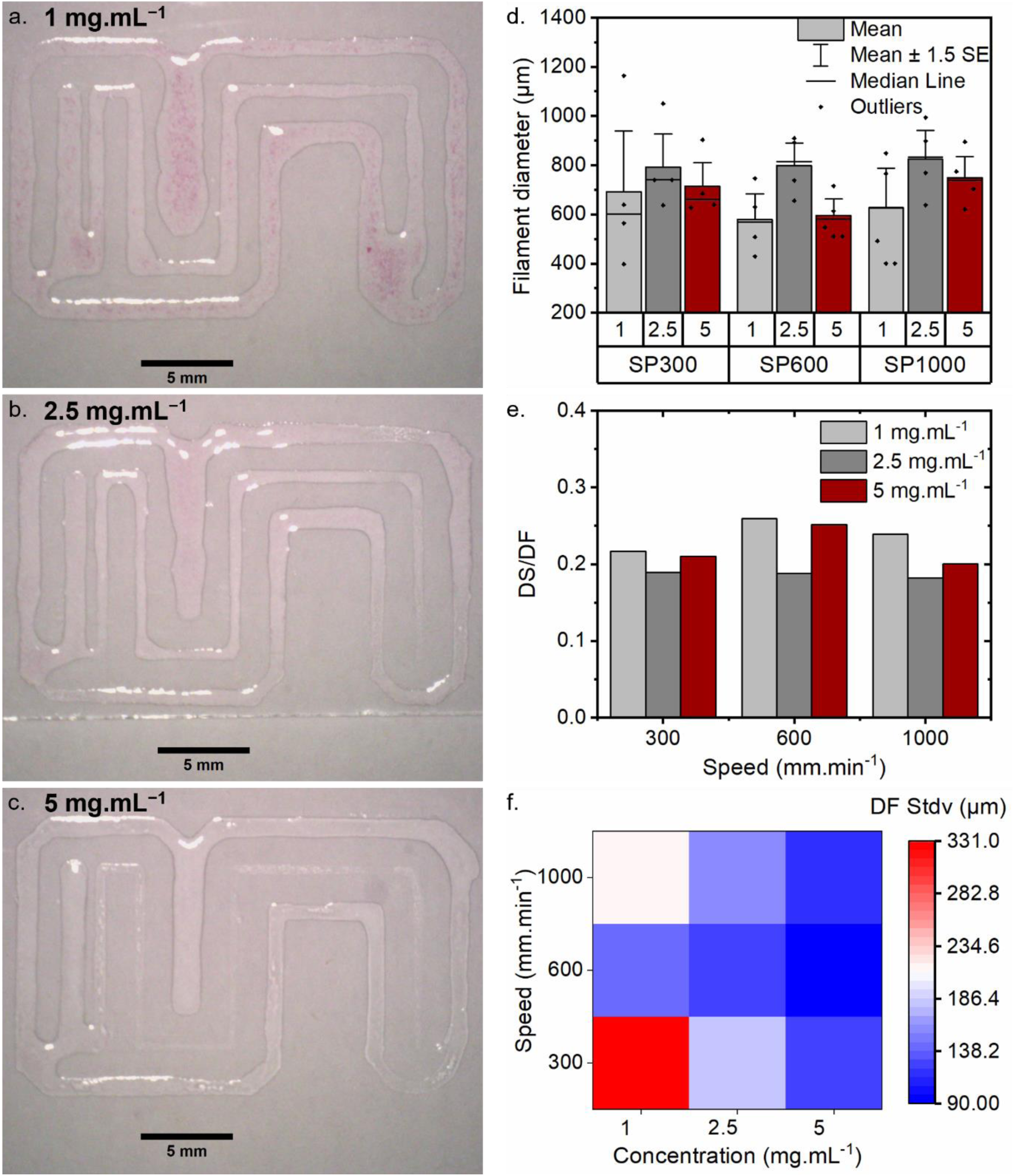
Printability of dECM bioinks on glass substrate. Pictures taken directly after printing at 600 mm.min ^−1^ with pre-gel at a. 1 mg.mL^−1^, b. 2.5 mg.mL^−1^ and c. 5 mg.mL^−1^, d. Diameter of the filament measured on the pictures, e. Ratio between the diameter of the syringe and the diameter of the filament DS/DF for all concentrations and speeds, f. heat map representing the standard deviation to the mean of filament diameter after printing.

In a second study, to reduce spreading of the bioink, the same serpentine pattern was printed with the ink at the concentration of 5 and 2.5 mg.mL^−1^ at 600 mm.min^−1^, on a polystyrene Petri dish (Fig. 5). The Petri dish is a relevant substrate as it is commonly used in cell culture, and 600 mm.min^−1^ was chosen as it was the speed leading to more homogeneous prints. Pictures of the prints are presented in Fig. 5a (2.5 mg.mL^−1^) and Fig. 5b (5 mg.mL^−1^). For both concentrations, the shape of the serpentine was preserved alongside the pattern. Indeed, the filament did not fuse, even for the smaller gap of 1 mm. The diameter of the filament was measured with the same method as for the glass substrate, and is presented in Fig. 5c. The width was between 420 and 540 µm, which is lower than on the glass slide, as anticipated. There was no statistical difference between the prints at 2.5 and 5 mg.mL^−1^. However, as observed on the glass substrate, the values are more dispersed with the pre-gel at 2.5 mg.mL^−1^ compared to the one at 5 mg.mL^−1^.

**Fig. 5.**
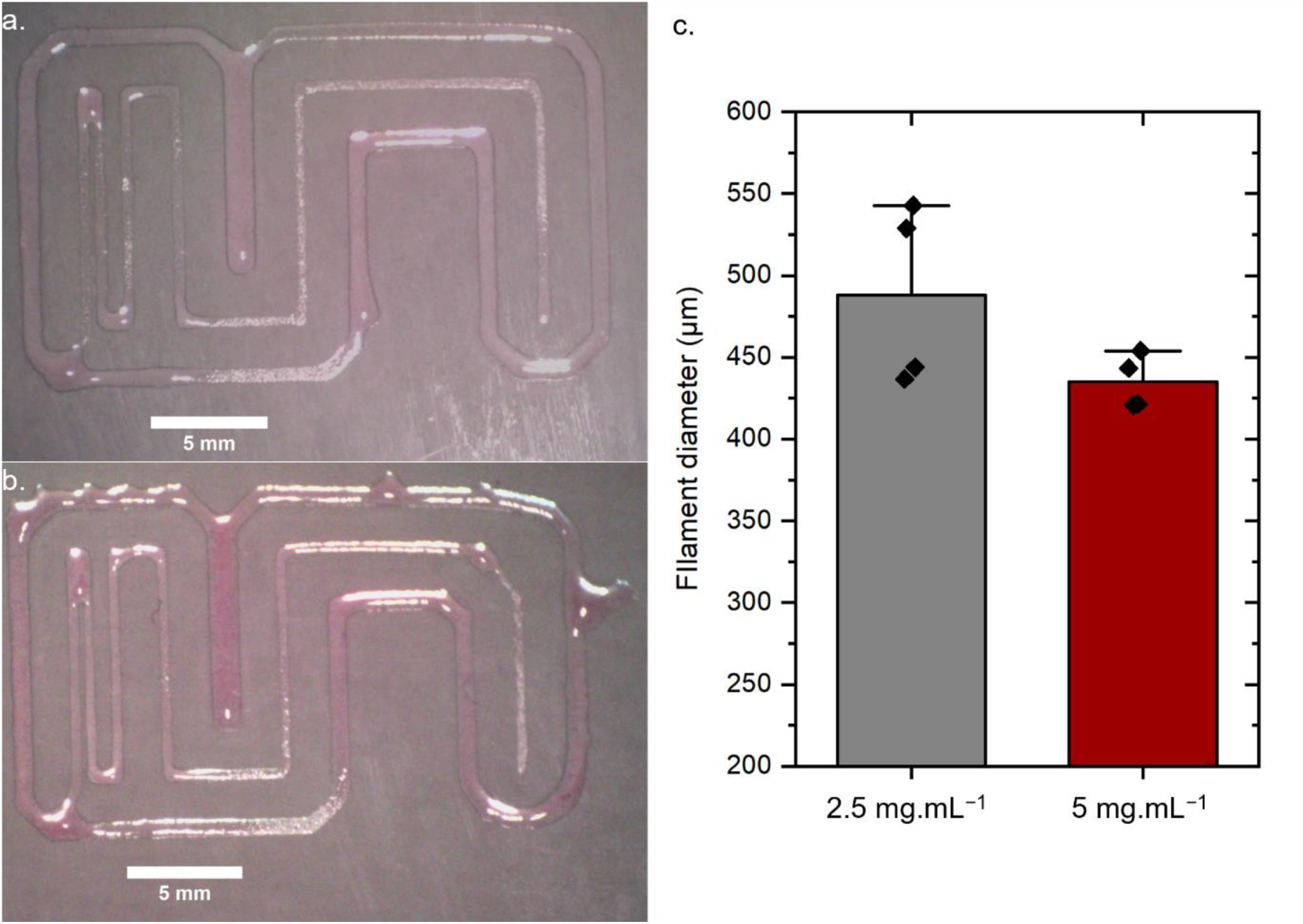
Printability of dECM bioinks on plastic substrate. Pictures taken directly after printing with pre-gel at a. 2.5 mg.mL^−1^ and b. 5 mg.mL^−1^, c. Diameter of the filament measured on the pictures.

In a third study, we aimed to print a grid pattern with several layers, using the pre-gel at 5 mg.mL^−1^, on plastic substrate, and with a speed of 600 mm.min^−1^. Fig. 6a shows a picture taken directly after printing of the grid pattern with two layers, and Fig. 6b shows the same pattern but with 4 layers. With only 2 layers, the pattern is preserved, with most of the empty areas preserved. On the contrary, it was not possible to print more than 4 layers due to collapsing of the construct.

**Fig. 6.**
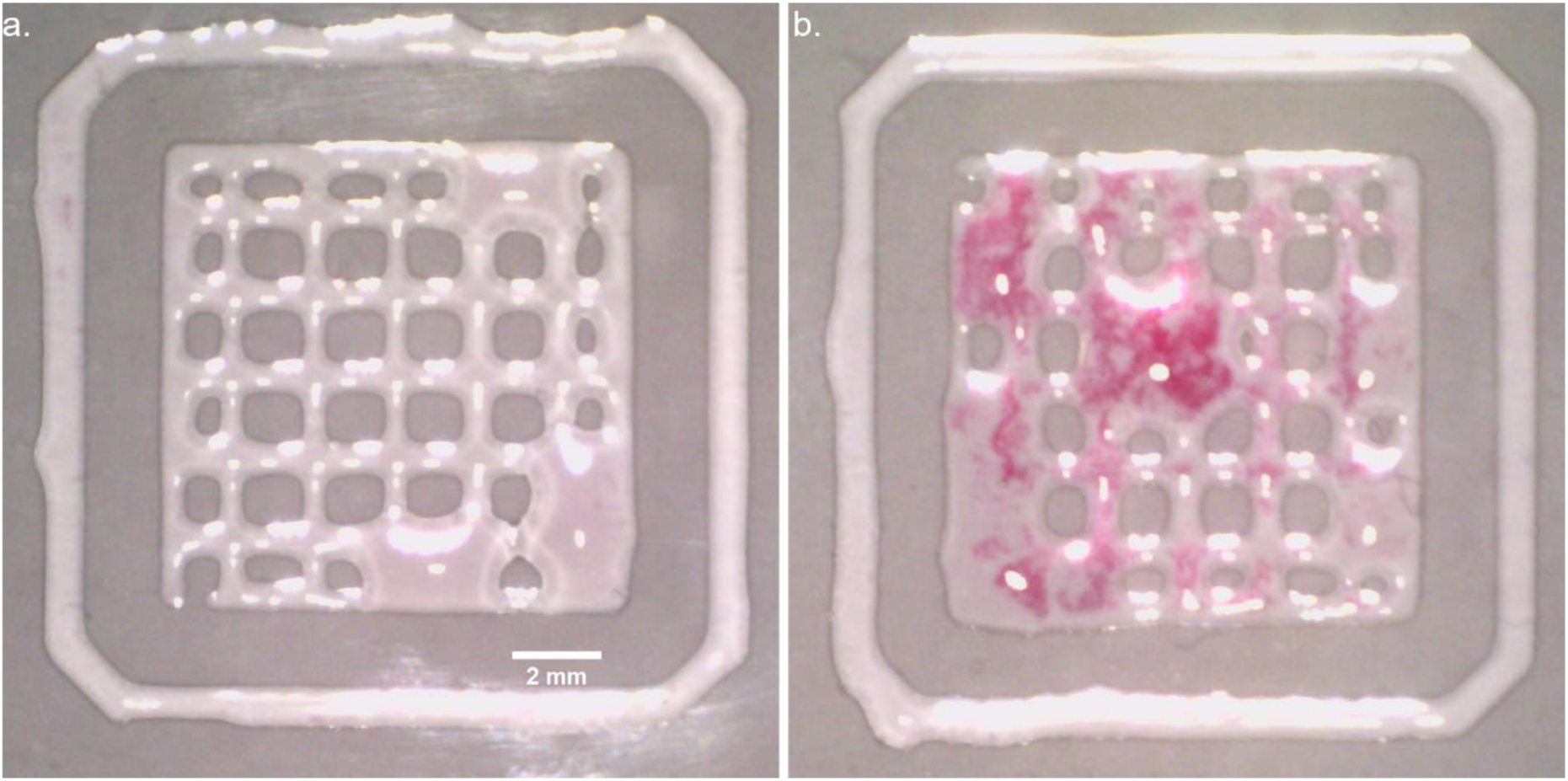
Layer stacking of printed dECM hydrogels. Pictures taken directly after printing with pre-gel at 5 mg.mL^−1^ at 600 mm.min ^−1^. a. 2 layers and b. 4 layers.

Overall, we showed extrudability and printability of the dECM pre-gels as ink for all concentrations. Prints are more homogeneous when using a more concentrated ink compared to a low-concentrated ink. The spreading of the filament was comparable with previous studies using dECM from porcine epidermidis^28^. At 5 mg.mL^−1^ without any further printing development it was possible to print one and two layers, after which the layer stacking becomes difficult. As previously observed with dECM inks, further printing development is needed to be able to print 3D structures. This could involve optimization of printing speed, needle size, substrate adhesion, dwell time between printed layers, use of supported bath, etc.

### Cell culture on dECM hydrogels

In the first instance, the cytocompatibility of dECM hydrogels was tested with fibroblast cells seeded on top of precast gels, at the three concentrations. dECM hydrogels were cast a day before seeding, and cell proliferation was monitored up to 7 days, with media renewal after 3 days. Cell proliferation was observed by confocal imaging after 24 hours, 72 hours and 7 days of culture.

With the hydrogel at 1 mg.mL^−1^, cells tend to sink down to the bottom of the plate as observed in Fig. 7. After 24 h of culture (Fig. 7a-c), two populations of cells are observed: at the bottom and within the gel. Cells at the bottom spread with a similar morphology to the control cells on plastic dishes. On the contrary, the cells present within the gel were mainly round shaped, indicating that they have not started to spread. After 72 h of culture (Fig. 7d-f), the cells within the hydrogels were elongated, with a morphology different than on the plastic control, indicating attachment to the gel and spreading. The cells that spread at the bottom of the plate proliferated and almost covered the entire surface. After 7 days of culture (Fig. 7g-i), the bottom of the well was covered by cells, and elongated cells were also visible in the gel. At 5 mg.mL^−1^, the cells stayed on top of the hydrogel for the entire duration of the experiment (Fig. 8). After 24 hours (Fig. 8a-b), they were round shaped. After 72 h, most cells were still round shaped (Fig. 8c-d), but they started to spread after 7 days, indicating their adherence to the hydrogel’s matrix (Fig. 8e-f). However, we observed fewer cells compared to cells grown on the plastic as a control, and in most cases, they were still round. With the hydrogel at 2.5 mg.mL^−1^, the cell behavior was intermediate to the ones observed at 5 and 1 mg.mL^−1^ (Fig. 9). After 24 h of culture, cells were on top of the gel and round-shaped (Fig. 9a-b). However, some of them started to elongate and penetrate the hydrogels after 72 h of culture (Fig. 9c-d). After 7 days, cells had penetrated the 3D environment of the gel and spread (Fig. 9e-f). In summary, cell response varies with dECM concentration in hydrogels. At high concentration, here 5 mg.mL^−1^, fibroblasts tended to spread less on the dECM hydrogels, while they proliferated on and within the gel at lower concentration, here 2.5 and 1 mg.mL^−1^.

**Fig. 7.**
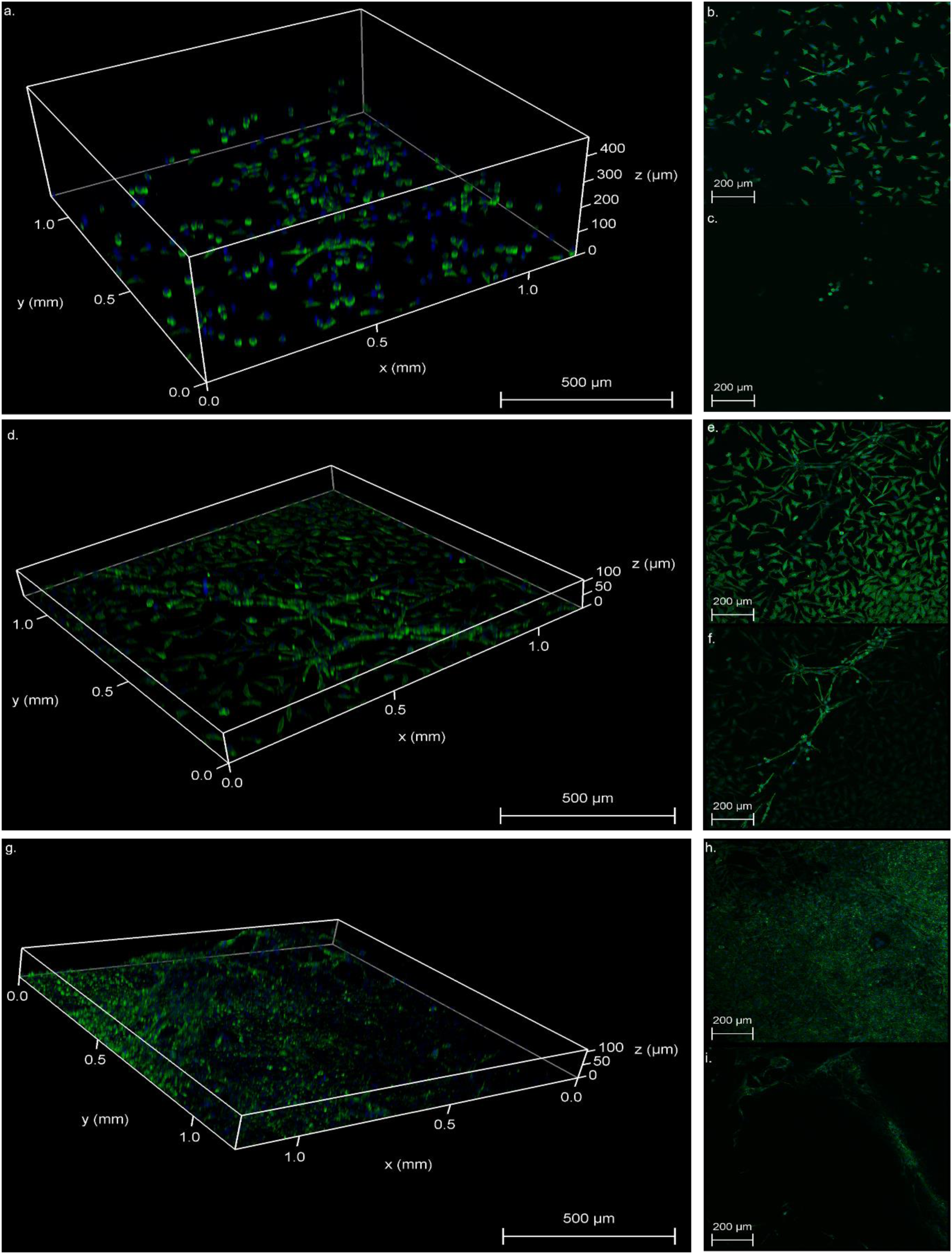
3D projection of confocal microscope imaging of L929 fibroblasts cultivated on top of 1 mg.mL^−1^ dECM hydrogel. a-c after 24 h of culture, d-f after 72 h of culture, and g-i after 7 days of culture. Panels b., e. and h. shows the bottom of the well, while c., f. and i. shows images taken in the middle of the gel.

**Fig. 8.**
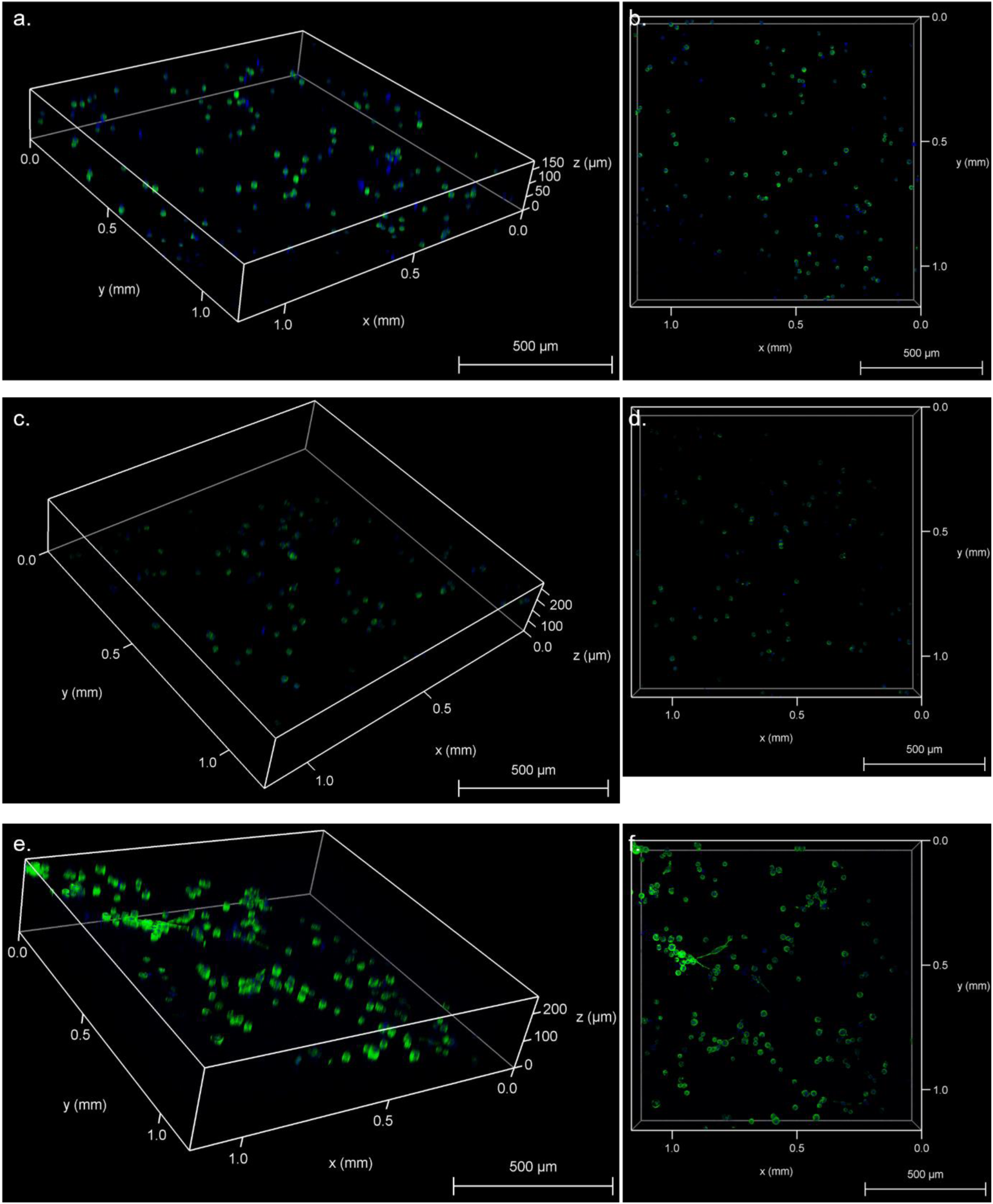
3D projection of confocal microscope imaging of L929 fibroblasts cultivated on top of 5 mg.mL^−1^ dECM hydrogel. a,b. after 24 h of culture, c,d. after 72 h of culture, and e,f. after 7 days of culture. Panels b., d., and f. show a top view.

**Fig. 9.**
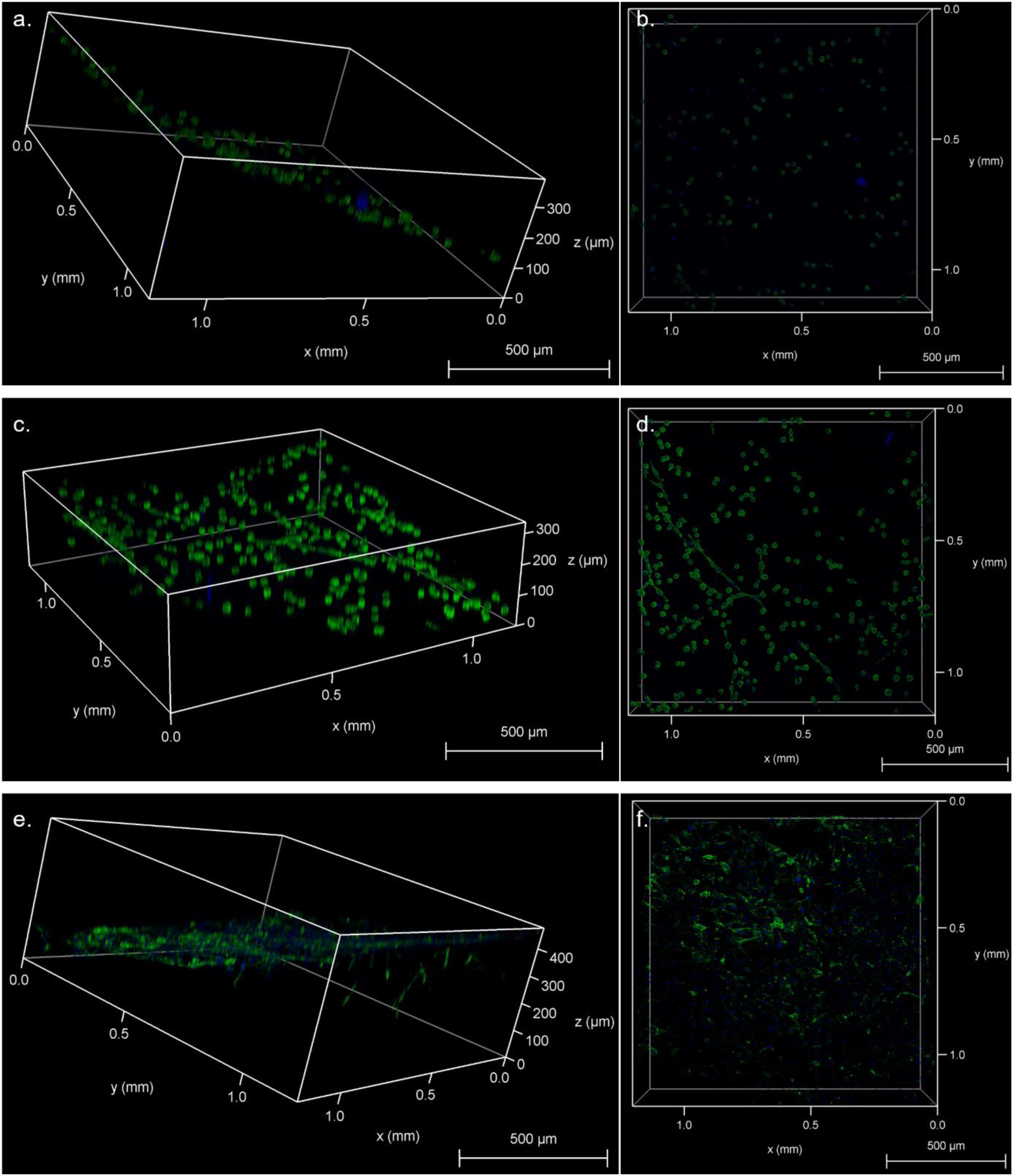
3D projection of confocal microscope imaging of L929 fibroblasts cultivated on top of 2.5 mg.mL^−1^ dECM hydrogel. a,b. after 24 h of culture, c,d. after 72 h of culture, and e,f. after 7 days of culture. Panels b., d., and f. show a top view.

The metabolic activity of the cells cultivated on top of the dECM hydrogels were studied with AlamarBlue assays, after 24 h, 72 h and 7 days, and are presented in Fig. S4a, with the results of the statistical analysis (one-way ANOVA followed by Tukey post-hoc test) presented in Fig. S4b. As the cells grow at different rates on the tested gels, it is difficult to directly compare the metabolic activity of the cells using AlamarBlue, because the assay depends on the number of the cells. However, the results serve to confirm the observations with confocal microscopy. At 24 h, the metabolic activity was the same for the cells cultivated on the three hydrogels. After 72 h of culture, the metabolic activity of the cells with the 1 mg.mL^−1^ concentration was significantly higher than the one for the cells with the hydrogels at 5 mg.mL^−1^, but there was no significant difference between the gels at 1 and 2.5 mg.mL^−1^ concentration and between the gels at 2.5 and 5 mg.mL^−1^ concentration. At 7 days, the metabolic activity was significantly higher with the hydrogel at 1 mg.mL^−1^ compared to the other gels, and at 2.5 mg.mL^−1^ compared to 5 mg.mL^−1^. At 72 h, the metabolic activity was similar to the one at 24 h, for all concentrations. However, it increased significantly after 7 days of culture, to different extents depending on the concentration.

The higher AlamarBlue reduction with the hydrogel at 1 mg.mL^−1^ can be related to the higher number of cells present in this case, from 24 h, as observed on the confocal images. Indeed, as cells are also largely present at the bottom of the well, they tend to proliferate faster. It is not surprising that the cells with the 5 mg.mL^−1^ hydrogel express a lower AlamarBlue reduction than with other hydrogel. As observed with confocal imaging, a lower number of cells was present with the 5 mg.mL^−1^ hydrogel, compared to other gels. Moreover, they were mainly round shaped, and it is known that round shaped cells have a lower metabolic activity than cells that have attached and spread. However, as the AlamarBlue reduction is significantly higher at 7 days, compared to 24 h and 72 h time points, the assay confirms that cells seeded on top of 5 mg.mL^−1^ hydrogel has started to proliferate after 7 days.

These findings show first that fibroblasts can be cultivated with skin-derived dECM hydrogel, even with the hydrogels at 5 mg.mL^−1^. However, within the first 24 h to 72 h the cells are less proliferative when seeded on top of the hydrogels at 5 and 2.5 mg.mL^−1^, as indicated by their round morphology and lower metabolic activity. This can be attributed to a change of pH or osmotic pressure when the hydrogel swells in the culture media.^21^ However, the cells accommodated for these difficult conditions and started to elongate and proliferate after 24 h, as also previously observed with dECM bioinks of various origin (tendon^34^, kidney^36^).

Our second finding is that the stiffness of the hydrogel will influence the cell migration within the gel’s matrix structure, as expected for this type of gels.^27^ For soft hydrogels, i.e. at 1 mg.mL^−1^ here, fibroblasts tend to directly sink down to the bottom of the well and adhere to the plastic. On the contrary, when the hydrogel is stiff, i.e. at 5 mg.mL^−1^ here, they cannot migrate as easily within the 3D environment and stay and proliferate to some extent on top of the gel. However, an intermediate stiffness allows for the penetration of the 3D environment of the gel, i.e. at 2.5 mg.mL^−1^ herein. The difference in migration can also be associated with the higher possibility for the cells to remodel the matrix in softer hydrogels, as well as the increased pore size within less concentrated hydrogels. Remodeling of the matrix is one of the migration mechanisms in cell growth in 3D matrixes.^37–40^ In addition, migrating cells within our dECM hydrogels elongated with a morphology different to the one observed on the plastic substrate. When grown on 2D rigid plastic substrates, cells tend to be flat with remodeling of the internal cytoskeleton.^38^ On the contrary, when grown in 3D environment, cells tend to organize into tissue-like structures. Moreover, in ECM mimicking-matrices, fibroblasts tend to align to the fibers and remodel the hydrogel matrix^37,38,41^, as seen in this work.

In summary, results for printability and cell proliferation and growth showed the importance to balance mechanical and biological properties when formulating bioinks. Indeed, cells cannot penetrate the 3D structure of the 5 mg.mL^−1^ gels, but they can proliferate within the gels at lower concentrations. The good printability and shape fidelity of the ink at 5 mg.mL^−1^ makes it suitable as a substrate for cell culture, with complex 2D shapes that can be achieved though extrusion-based 3D printing. However, as the cells cannot penetrate the 3D environment of the hydrogel, this ink cannot be used as bioink for 3D bioprinting. At 1 mg.mL^−1^, the gelation kinetics are very slow and the final stiffness very low. Moreover, the shape fidelity after printing is not sufficient as the prints are very heterogeneous. Even though the gels are not cytotoxic to the cells, and cells can thrive in presence of the gel, they do not adhere sufficiently to the gel and sink down directly to the bottom of the plate. Overall, at 1 mg.mL^−1^, the performance of the ink is deemed insufficient to be used as an ink or bioink. The ink at 2.5 mg.mL^−1^ is a good compromise between these two extremes, with possible use as a bioink. Indeed, the intermediate stiffness of the hydrogel (100 Pa) allows the cells to penetrate the 3D environment and spread within the gel. On plastic substrates, the ink can be printed and retain the shape of the design. However, the gelation kinetics of the gel is lower than at 5 mg.mL^−1^. To make the most of this ink as bioink and to create complex 3D structures, the use of a support bath^36^ or support material^11^ could be advantageous, as is commonly done with other dECM inks. Another possibility to improve the printability of tissue-derived inks is to print with the hydrogel after gelation, as previously described in the literature,^34^ using the shear-thinning properties of dECM hydrogels.

### Conclusions

Skin-derived inks with tunable gelation kinetics, rheological properties, printability and cytocompatibility were successfully developed. With a variation of concentration, we were able to obtain hydrogels with storage modulus ranging from 10 Pa to more than 1 kPa, with gelation kinetics ranging from few minutes to several hours at 37 °C. The dECM inks were printable with extrusion-based 3D printing, even though the ink spread extensively on a glass substrate. Our printability study shows that the inks at 2.5 and 5 mg.mL^−1^ are more suitable for printing than the 1 mg.mL^−1^ ink, with better shape fidelity for the ink at 5 mg.mL^−1^. Results for cell proliferation and growth within the hydrogels highlights the importance of mechanical properties in regulating cell migration. When grown on top of the softer hydrogels (1 mg.mL^−1^ and 2.5 mg.mL^−1^), fibroblasts cells spread and infiltrated the 3D matrix of the hydrogels, while they remained on top of the stiffer hydrogels (5 mg.mL^−1^). Our findings demonstrate the potential of porcine skin-derived hydrogels with tunable properties for 3D bioprinting applications, enabling fast and reproducible fabrication of dECM environments. This opens the door for further studies in tissue engineering or in vitro cellular investigations.

## Author contributions

Conceptualization: EP, MA, and CP, data curation: EP, AMM, IB, formal analysis: EP, AMM, IB, MA, CP, funding acquisition: MA and CP, investigation: EP, AMM, IB, methodology: EP, AMM, MA and CP, project administration: MA and CP, resources: MA and CP, supervision: MA and CP, writing – original draft: EP, MA, CP, and writing – review & editing: all authors

## Data availability

Data are available upon request.

## Acknowledgements

We thank the funding support of Carl Tryggers Stiftelse (CTS 22:2367). We thank our lab members Karl Vilhem Ståhl and August Claesson for assistance during experiments. We thank Oliver Donzel-Gargand for his help with SEM imaging, and Myfab Uppsala for providing facilities. Myfab is funded by the Swedish Research Council (2019-00207) as a national research infrastructure. Sergio Estrada, Veronika Wingstedt and The Preclinical PET-MRI Platform of Uppsala University are acknowledged for access and assistance with microtomy. This work was conducted within the Additive Manufacturing for the Life Sciences Competence Center (AM4Life). The authors acknowledge financial support from Sweden’s Innovation Agency VINNOVA (Grant no: 2019-00029).

## References

1 Information about Organ, Eye, and Tissue Donation | organdonor.gov, https://www.organdonor.gov/, (accessed October 1, 2024).

2 A. N. Carrier, A. Verma, M. Mohiuddin, M. Pascual, Y. D. Muller, A. Longchamp, C. Bhati, L. H. Buhler, D. G. Maluf and R. P. H. Meier, Front. Immunol., DOI:10.3389/fimmu.2022.900594.

3 R. Xie, V. Pal, Y. Yu, X. Lu, M. Gao, S. Liang, M. Huang, W. Peng and I. T. Ozbolat, Biomaterials, 2024, 304, 122408.

4 W. M. S. Russell, R. L. Burch and C. W. Hume, The principles of humane experimental technique, Methuen London, 1959, vol. 238.

5 W. He, J. Deng, B. Ma, K. Tao, Z. Zhang, S. Ramakrishna, W. Yuan and T. Ye, ACS Appl. Bio Mater., 2024, 7, 17–43.

6 S. Chae and D.-W. Cho, Acta Biomater., 2023, 156, 4–20.

7 A. Lee, A. R. Hudson, D. J. Shiwarski, J. W. Tashman, T. J. Hinton, S. Yerneni, J. M. Bliley, P. G. Campbell and A. W. Feinberg, Science, 2019, 365, 482–487.

8 R. Khoeini, H. Nosrati, A. Akbarzadeh, A. Eftekhari, T. Kavetskyy, R. Khalilov, E. Ahmadian, A. Nasibova, P. Datta, L. Roshangar, D. C. Deluca, S. Davaran, M. Cucchiarini and I. T. Ozbolat, Adv. NanoBiomed Res., 2021, 1, 2000097.

9 A. Behre, J. W. Tashman, C. Dikyol, D. J. Shiwarski, R. J. Crum, S. A. Johnson, R. Kommeri, G. S. Hussey, S. F. Badylak and A. W. Feinberg, Adv. Healthc. Mater., 2022, 11, 2200866.

10 C. Mandrycky, Z. Wang, K. Kim and D.-H. Kim, Biotechnol. Adv., 2016, 34, 422–434.

11 F. Pati, J. Jang, D.-H. Ha, S. Won Kim, J.-W. Rhie, J.-H. Shim, D.-H. Kim and D.-W. Cho, Nat. Commun., 2014, 5, 3935.

12 Y. S. Zhang, G. Haghiashtiani, T. Hübscher, D. J. Kelly, J. M. Lee, M. Lutolf, M. C. McAlpine, W. Y. Yeong, M. Zenobi-Wong and J. Malda, Nat. Rev. Methods Primer, 2021, 1, 75.

13 Z. Fu, S. Naghieh, C. Xu, C. Wang, W. Sun and X. Chen, Biofabrication, 2021, 13, 033001.

14 M. Zhe, X. Wu, P. Yu, J. Xu, M. Liu, G. Yang, Z. Xiang, F. Xing and U. Ritz, Materials, 2023, 16, 3197.

15 J. A. DeQuach, V. Mezzano, A. Miglani, S. Lange, G. M. Keller, F. Sheikh and K. L. Christman, PLOS ONE, 2010, 5, e13039.

16 T. L. Sellaro, A. K. Ravindra, D. B. Stolz and S. F. Badylak, Tissue Eng., 2007, 13, 2301–2310.

17 H. Zhang, Y. Wang, Z. Zheng, X. Wei, L. Chen, Y. Wu, W. Huang and L. Yang, Theranostics, 2023, 13, 2562–2587.

18 D. Kim and G. Kim, Biofabrication, 2023, 15, 045006.

19 M. Ali, A. K. Pr, J. J. Yoo, F. Zahran, A. Atala and S. J. Lee, Adv. Healthc. Mater., 2019, 8, 1800992.

20 R. Naomi, P. M. Ridzuan and H. Bahari, Polymers, 2021, 13, 2642.

21 F. Zhao, J. Cheng, J. Zhang, H. Yu, W. Dai, W. Yan, M. Sun, G. Ding, Q. Li, Q. Meng, Q. Liu, X. Duan, X. Hu and Y. Ao, Acta Biomater., 2021, 131, 262–275.

22 S. Y. Chun, Y.-S. Ha, B. H. Yoon, E. H. Lee, B. M. Kim, H. Gil, M.-H. Han, T. G. Kwon, B. S. Kim and J. N. Lee, J. Biomed. Mater. Res. A, 2022, 110, 928–942.

23 V. Coulson-Thomas and T. Gesteira, BIO-Protoc., DOI:10.21769/BioProtoc.1236.

24 S. L. Voytik-Harbin, A. O. Brightman, B. Z. Waisner, J. P. Robinson and C. H. Lamar, Tissue Eng., 1998, 4, 157–174.

25 A. Engberg, C. Stelzl, O. Eriksson, P. O’Callaghan and J. Kreuger, Sci. Rep., 2021, 11, 21547.

26 S. Girardeau-Hubert, B. Lynch, F. Zuttion, R. Label, C. Rayee, S. Brizion, S. Ricois, A. Martinez, E. Park, C. Kim, P. A. Marinho, J.-H. Shim, S. Jin, M. Rielland and J. Soeur, Acta Biomater., 2022, 143, 100–114.

27 M. T. Wolf, K. A. Daly, E. P. Brennan-Pierce, S. A. Johnson, C. A. Carruthers, A. D’Amore, S. P. Nagarkar, S. S. Velankar and S. F. Badylak, Biomaterials, 2012, 33, 7028–7038.

28 G. Ahn, K.-H. Min, C. Kim, J.-S. Lee, D. Kang, J.-Y. Won, D.-W. Cho, J.-Y. Kim, S. Jin, W.-S. Yun and J.-H. Shim, Sci. Rep., 2017, 7, 8624.

29 J. Karvinen and M. Kellomäki, Bioprinting, 2023, 32, e00274.

30 H. Kim, M.-N. Park, J. Kim, J. Jang, H.-K. Kim and D.-W. Cho, J. Tissue Eng., 2019, 10, 2041731418823382.

31 Y. Yang and L. J. Kaufman, Biophys. J., 2009, 96, 1566–1585.

32 B. Wang, X. Barceló, S. Von Euw and D. J. Kelly, Mater. Today Bio, 2023, 20, 100624.

33 J. Wu, L. Fu, Z. Yan, Y. Yang, H. Yin, P. Li, X. Yuan, Z. Ding, T. Kang, Z. Tian, Z. Liao, G. Tian, C. Ning, Y. Li, X. Sui, M. Chen, S. Liu and Q. Guo, Biomater. Res., 2023, 27, 7.

34 B. Toprakhisar, A. Nadernezhad, E. Bakirci, N. Khani, G. A. Skvortsov and B. Koc, Macromol. Biosci., 2018, 18, 1800024.

35 L. T. Saldin, M. C. Cramer, S. S. Velankar, L. J. White and S. F. Badylak, Acta Biomater., 2017, 49, 1–15.

36 R. Sobreiro-Almeida, M. Gómez-Florit, R. Quinteira, R. L. Reis, M. E. Gomes and N. M. Neves, Biofabrication, 2021, 13, 045006.

37 S. Rhee and F. Grinnell, Adv. Drug Deliv. Rev., 2007, 59, 1299–1305.

38 J. P. Woodley, D. W. Lambert and I. O. Asencio, Tissue Eng. Part B Rev., 2022, 28, 569–578.

39 J. S. Harunaga and K. M. Yamada, Matrix Biol., 2011, 30, 363–368.

40 K. M. Yamada and M. Sixt, Nat. Rev. Mol. Cell Biol., 2019, 20, 738–752.

41 K. M. Hakkinen, J. S. Harunaga, A. D. Doyle and K. M. Yamada, Tissue Eng. Part A, 2011, 17, 713–724.

