## Supplementary Information for "Tuning the Mechanical Properties and Printability of Viscoelastic Skin-Derived Hydrogels for 3D Cell Culture"

**Fig. S1** Quantification of the molecular weight of collagen chains in porcine skin-derived dECM gels at 5 mg.ml^−1^ concentration, assessed with SDS Page assay.

**Fig. S2** Gelation kinetics. Representative time sweep curves at a. 5 mg.mL^−1^, b. 2.5 mg.mL^−1^, c. 1 mg.mL^−1^. Evolution of phase angle and axial force during gelation kinetics at d. 5 mg.mL^−1^, e. 2.5 mg.mL^−1^

**Fig. S3** Pictures of print on glass slides, all speed and all concentrations

**Fig. S4** Metabolic activity of L929 fibroblasts cultivated on dECM gels over 7 days.

**Table 1** Statistics performed with one-way ANOVA followed by Tukey post hoc test: * represents p < 0.05, ** represents p < 0.01 and *** represents p < 0.001

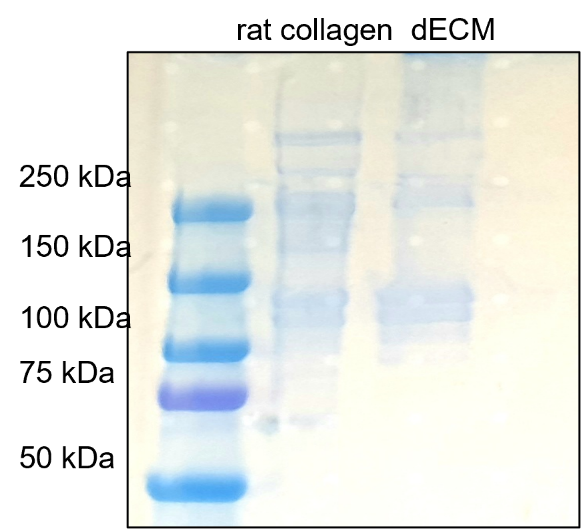

**Fig. S1** Quantification of the molecular weight of collagen chains in porcine skin-derived dECM gels at 5 mg.ml^−1^ concentration, assessed with SDS Page assay.

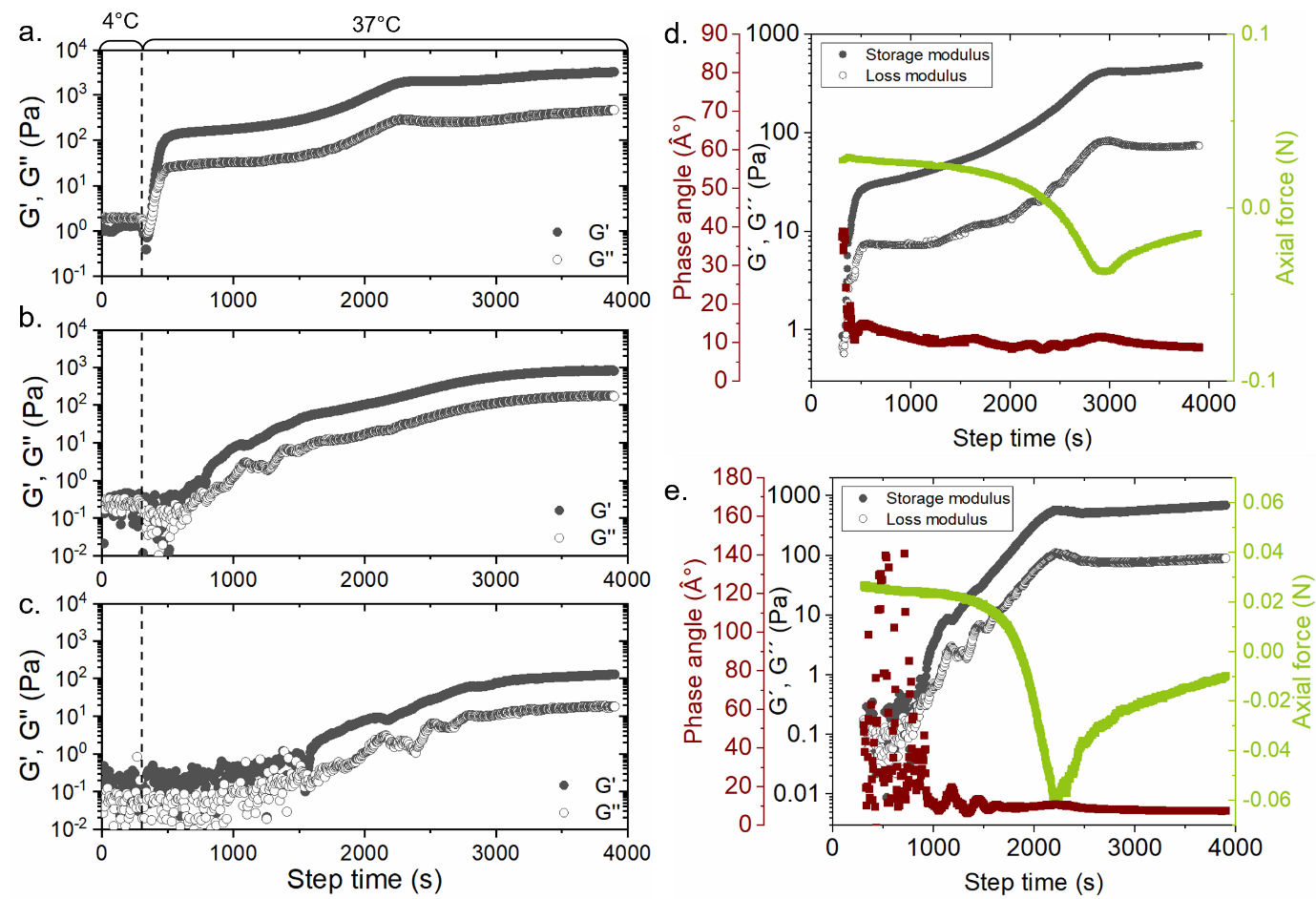

**Fig. S2** Gelation kinetics. Representative time sweep curves at a. 5 mg.mL^−1^, b. 2.5 mg.mL^−1^, c. 1 mg.mL^−1^. Evolution of phase angle and axial force during gelation kinetics at d. 5 mg.mL^−1^, e. 2.5 mg.mL^−1^

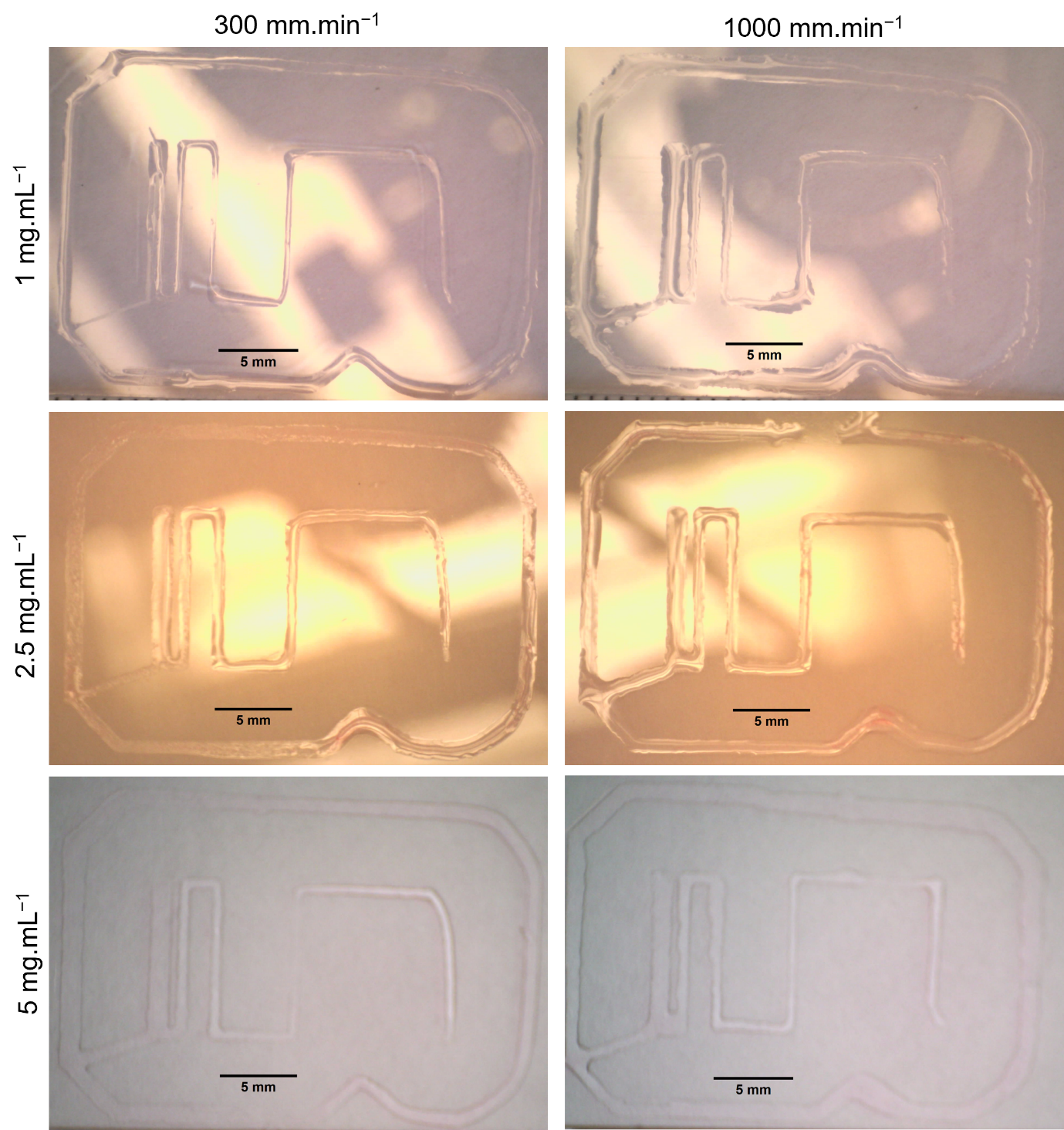
 **Fig. S3** Pictures of print on glass slides, at 300 and 1000 mm.min^−1^, for all dECM concentrations.

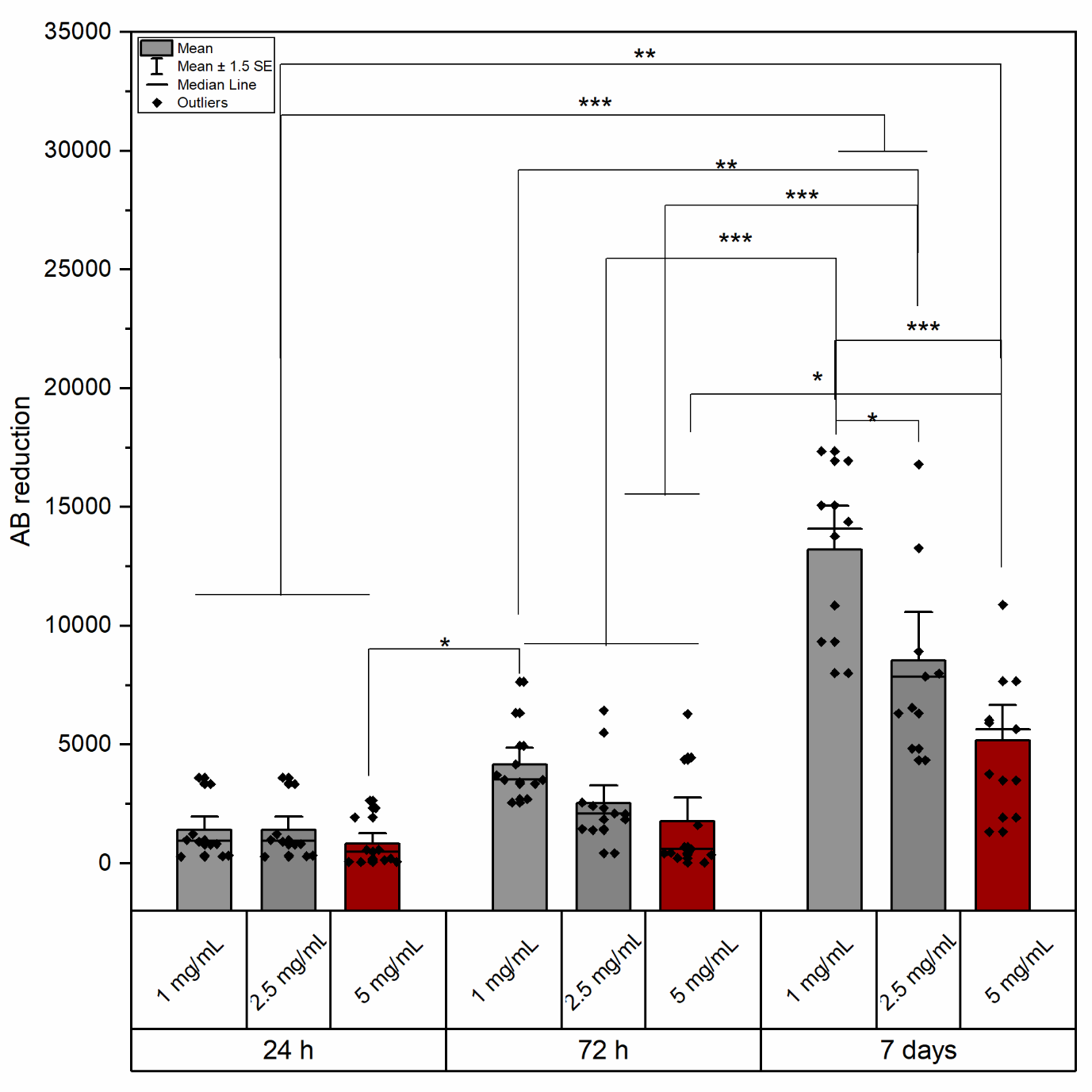

**Fig. S4** Metabolic activity of L929 fibroblasts cultivated on dECM gels over 7 days. * Represents p < 0.05, ** represents p < 0.01 and *** represents p < 0.001

|  | 24 h - 1 mg/mL | 24 h - 2.5 mg/mL | 24 h - 5 mg/mL | 72 h - 1 mg/mL | 72 h - 2.5 mg/mL | 72 h - 5 mg/mL | 7 d - 1 mg/mL | 7 d - 2.5 mg/mL | 7 d - 5 mg/mL |
| --- | --- | --- | --- | --- | --- | --- | --- | --- | --- |
| 24 h - 1 mg/mL |  | 1 | 0.99938 | 0.09549 | 0.95056 | 0.99999 | < 0.0001 | < 0.0001 | 0.00783 |
| 24 h - 2.5 mg/mL |  |  | 0.99938 | 0.09549 | 0.95056 | 0.99999 | < 0.0001 | < 0.0001 | 0.00783 |
| 24 h - 5 mg/mL |  |  |  | 0.02093 | 0.66819 | 0.98604 | < 0.0001 | < 0.0001 | 0.00135 |
| 72 h - 1 mg/mL |  |  |  |  | 0.71953 | 0.24709 | < 0.0001 | 0.00129 | 0.9842 |
| 72 h - 2.5 mg/mL |  |  |  |  |  | 0.99624 | < 0.0001 | < 0.0001 | 0.17237 |
| 72 h - 5 mg/mL |  |  |  |  |  |  | < 0.0001 | < 0.0001 | 0.02883 |
| 7 d - 1 mg/mL |  |  |  |  |  |  |  | 0.00157 | < 0.0001 |
| 7 d - 2.5 mg/mL |  |  |  |  |  |  |  |  | 0.05234 |
| 7 d - 5 mg/mL |  |  |  |  |  |  |  |  |  |

**Table 1** Statistics performed with one-way ANOVA followed by Tukey post hoc test. Table shows p values.
